# rMVP: A Memory-efficient, Visualization-enhanced, and Parallel-accelerated tool for Genome-Wide Association Study

**DOI:** 10.1101/2020.08.20.258491

**Authors:** Lilin Yin, Haohao Zhang, Zhenshuang Tang, Jingya Xu, Dong Yin, Zhiwu Zhang, Xiaohui Yuan, Mengjin Zhu, Shuhong Zhao, Xinyun Li, Xiaolei Liu

## Abstract

Along with the development of high-throughout sequencing technologies, both sample size and number of SNPs are increasing rapidly in Genome-Wide Association Studies (GWAS) and the associated computation is more challenging than ever. Here we present a Memory-efficient, Visualization-enhanced, and Parallel-accelerated R package called “rMVP” to address the need for improved GWAS computation. rMVP can: (1) effectively process large GWAS data; (2) rapidly evaluate population structure; (3) efficiently estimate variance components by EMMAX, FaST-LMM, and HE regression algorithms; (4) implement parallel-accelerated association tests of markers using GLM, MLM, and FarmCPU methods; (5) compute fast with a globally efficient design in the GWAS processes; and (6) generate various visualizations of GWAS related information. Accelerated by block matrix multiplication strategy and multiple threads, the association test methods embedded in rMVP are approximately 5-20 times faster than PLINK, GEMMA, and FarmCPU_pkg. rMVP is freely available at https://github.com/xiaolei-lab/rMVP.

## Introduction

The computation burden of GWAS is partially caused by the increasing sample size and marker density applied for these studies. As a result, how to efficiently analyse the big data is a big challenge. Additionally, GWAS have been widely used for detecting candidate genes that control human diseases and agricultural economic traits, where the accuracy of the results is of significant implications. Thus, how to achieve higher statistical power under a reasonable level of type I error is another challenge[1]. To efficiently detect more candidate genes with lower false positive rates is the current working goal for GWAS algorithms and tools[2, 3].

Introducing the population structure concept into GWAS has dramatically improved accuracy of detection. For example, incorporating the fractions of individuals belonging to subpopulations, namely Q matrix, reduces both false positive and false negative signals[4]. Principal components (PCs) are widely used to represent subpopulations and to enable the incorporation of population structure into GWAS[5]. Implementing the General Linear Model (GLM) to incorporate either the Q matrix or PCs as covariates, PLINK has become the most popular software package for GWAS[6].

False positives also stem from individuals that exhibit high variability in pairwise relatedness presumptively classified into different subpopulations. In addition to integrating population structure, statistical power can be substantially improved by the incorporation of hidden relationships in a mixed linear model (MLM) - particularly when population structure is less dominant than the cryptic relatedness[7]. Multiple algorithms have been developed to boost both the computational efficiency and statistical power of MLM methods[8–11]. Various software packages have also been developed for the implementation of these algorithms, including TASSEL[12], GAPIT[13, 14], GenABEL[15], EMMAX[16], GEMMA[17], and GCTA[18]. Even though the number of GWAS literature applying MLM-based methods is increasing rapidly, it is still not comparable to that of PLINK, primarily because PLINK operates much faster than MLM-based method software.

Besides the difference in computing time, MLM does not provide high statistical power compared to GLM. The difference in statistical power between GLM and MLM is negligible in some scenarios, such as mapping genes under the same false discovery rate in populations with strong population structure[19]. These populations include human populations, as well as animal and plant populations that have been isolated by breeding programs. Our newly developed method, FarmCPU (Fixed and random model Circulating Probability Unification) has higher statistical power than both GLM and MLM for evaluating populations with either weak or strong population structure. FarmCPU splits MLM into a fixed effect model (FEM) and a random effect model (REM), using them iteratively to increase the power for detecting candidate genes associated with population structure. Association tests in FarmCPU are validated by FEM with the same computing efficiency as GLM while the statistical power surpasses that of MLM at the same level of type I error.

Although recently developed methods have improved statistical power under certain assumptions, determining the most appropriate method for a given dataset is still convoluted. Human genetic studies often use large datasets with simple models, while plant and animal genetic studies prefer complex models with limited sample sizes. For a specific trait, it is usually difficult to identify the real genetic architecture and the most appropriate method to be used. Researchers have to try out multiple methods and identify candidate genes based on both statistical and biological evidence. Additionally, existing GWAS software rarely focuses on providing a flexible plotting function to display GWAS related information in a way that satisfies the personal aesthetic requirements of the researchers. Furthermore, with the development of multi-traits methods, such as GSA-SNP2[20], MTMM[21], mvLMMs[22], and mtSet[23], results from multiple-group GWAS need to be displayed simultaneously for easier comparisons. Therefore, there appears a need for analysing big data with limited computing memory, reasonable time, and reduced false positive rates, while displaying the results in high-quality figures. To address all of the above requirements, we developed the Memory-efficient, Visualization-enhanced, and Parallel-accelerated package (rMVP) in R.

## Methods and materials

We split the entire GWAS procedure into six sections: data preparation, evaluation of population structure, estimation of variance components, association tests, globally efficient design on GWAS process computing, and result visualization. Abundant functions have been implemented in rMVP for each section:

1. **Data preparation.** rMVP accepts multiple popular formats for genotype files, such as PLINK binary, Hapmap, VCF, and Numeric (e.g., genotype data can be coded as integer (0, 1, 2) or dosage/probability (0.1, 0.3, 0.6)). All above formats will be converted to the ‘big.matrix’ format. The advantage of converting genotype files into ‘big.matrix’ is that the size of the file is only limited by the storage capacity of the hard disk but not the processing capacity of Random Access Memory (RAM, and ‘memory’ is referred to RAM in this manuscript)[24].
2. **Evaluation of population structure and individual relationship.** For population structure analysis, PCs can be calculated using all available markers. An ideal population for GWAS assumed that the individuals were randomly selected from a big population, but the population could always be classified to multiple subpopulations in fact. The alleles with different frequencies in different subpopulations would generate false positives, we recommend to integrate the 3-5 top PCs as covariates into model to control false positives caused by population structure following previous studies[5, 19]. VanRaden Method is implemented in rMVP for the efficient construction of genomic relationship matrix (GRM)[25].
3. **Estimation of variance components.** Four algorithms are implemented for estimating variance components in rMVP: Brent (default method in rMVP)[26], EMMAX (Efficient Mixed-Model Association eXpedited) / P3D (Population Parameters Previously Determined)[8, 16], FaST-LMM (Factored Spectrally Transformed Linear Mixed Model) algorithms[9] and HE regression (Haseman-Elston regression)[27]. Different algorithms have different performances in terms of accuracy and efficiency. For instance, the Brent and EMMAX use eigen decomposition on genomic relationship matrix to avoid computing the inverse of big matrix, the FaST-LMM uses singular value decomposition on genotype matrix, which can be more efficient when the number of markers is far less than the number of individuals, the HE regression, which uses linear regression model to fit the similarity of phenotype and genomic relationship matrix among individuals, is less accurate but can be much more memory-efficient and time-saving, making it more promising in very large datasets.
4. **Association tests.** General Linear Model, Mixed Linear Model, and FarmCPU methods are implemented in rMVP for association tests. When there is more than one covariate (e.g. PCs) added to association test models, the inverse of the design matrix corresponding to the covariates will be calculated n times, where n is marker size. Block matrix multiplication strategy can be used to speed up the processes including inverse of the design matrix corresponding to the covariates and the testing markers. This strategy is used in all available association test methods in rMVP. Take GLM as an example, the fixed effect model function can be written as:

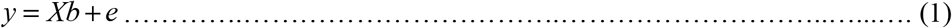

where *y* is a vector of phenotype, *X* is a matrix of fixed effects and test SNP, *b* is an incidence matrix for *X*, and *e* is a vector of residual that followed a normal distribution with mean of zero and 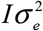 covariance, where *I* is the identity matrix and 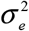 is the unknown residual variance. Equation (1) can be reformulated by following steps:

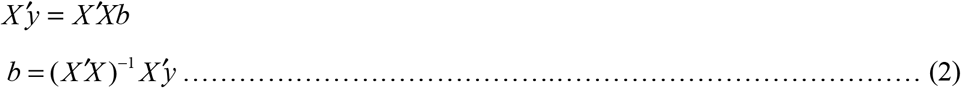 Where *X*′ is the transpose matrix of *X*. If there are k fixed effect vectors added as covariates in the model, *X* and *b* can be written as:

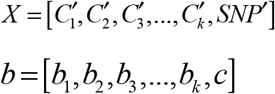

where *C*_1_, *C*_2_, *C*_3_,…,*C_k_* represent k fixed effect vectors and *SNP* represents the test SNP vector. Equation (2) can be written as

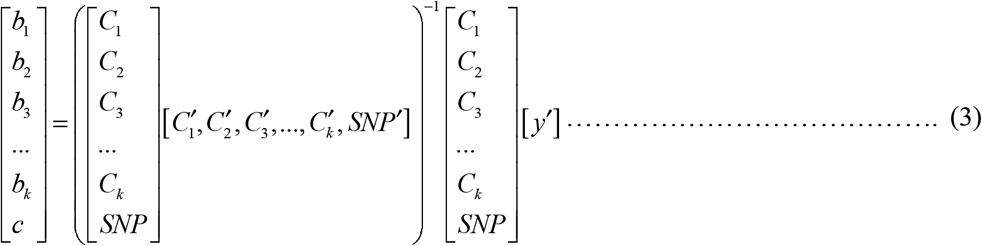 The most time-consuming part in equation (3) is the inverse of *M* matrix. And *M* is defined as:

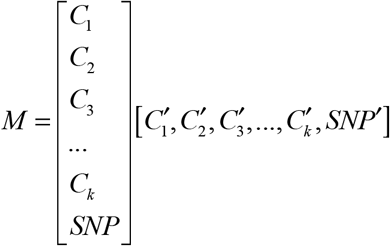 If we use *w* and *x* represent *C*_1_, *C*_2_, *C*_3_,…, *C_k_* and *SNP*, respectively, the inverse of *M* matrix can be written as:

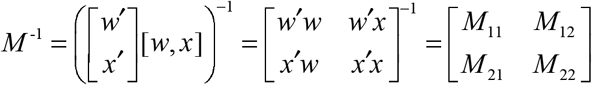

where

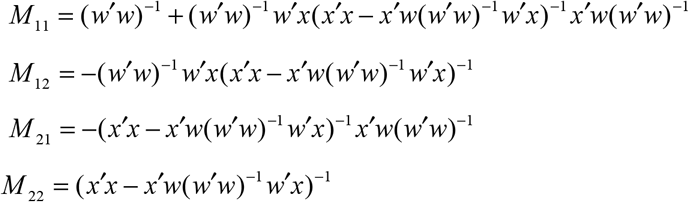 The inversion of *w*′*w* matrix is recomputed n times when constructing *M* _11_, *M* _12_, *M* _21_, *M* _22_ matrix for each test marker. For the matrix operations in GLM, MLM, and each iteration of FarmCPU, the *w* matrix is fixed, and the inversion of *w*′*w* can be calculated only once using block matrix multiplication strategy. As it is repeated n times when testing the SNPs, more time will be saved when there are more covariates in the model or more SNPs to be tested.
5. **Globally efficient design of GWAS calculations.** A standard GWAS pipeline generally includes PC derivation, GRM construction, variance components estimation, and association tests. There are three commonly used strategies for deriving the PCs: (a) the Eigen decomposition results of the matrix that represents the correlation coefficients between pairs of markers could be derived by (*M ^T^ M*)*v* = *λv*, where *M* is a *n* by *m* scaled genotype matrix (*n* is the number of individuals, *m* is the number of SNPs), the Eigen decomposition analysis is conducted on the correlation matrix *M ^T^ M*, the dimension of which is *m* by *m*, and this would lead to high requirements of both memory and computing time with the increasing number of SNPs; (b) The Singular Value Decomposition analysis could be conducted on the *M* matrix by *M* = *U*Σ*V*\*, its computational complexity is relative smaller than the method that described in (1), as it only needs to decompose a *n* by *m* matrix; (c) The PCs could be also derived by performing the Eigen decomposition of the GRM, which could be calculated by *GRM* = *M ^T^ M* / *m*, and its dimension is *n* by *n*. In the majority of cases, the number of markers (*m*) is far more than the number of individuals (*n*), this method has the smallest computational complexity compared with the other two. Moreover, the construction of GRM is always a key part in commonly used GWAS procedure, which has been precomputed already. Not only that, as shown in Figure S3, the Eigen decomposition results of GRM could be easily applied to processes of variance components estimation and association tests. By the default sets in rMVP, the Eigen decomposition analysis was conducted on GRM, which was constructed by VanRaden method[25], the methodological formula of VanRaden method can be defined as:

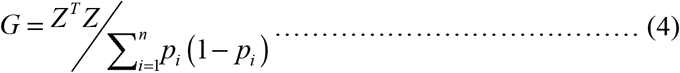 Where *Z* is a dimension of *m* by *n* matrix, *m* is the number of markers and *n* is the number of individuals, it can be derived from cantering the additive genotype matrix which was coded as 0, 1, 2 for genotype AA, AB, BB respectively, *p* is the minor allele frequency. After the eigen decomposition was finished, the eigen values and eigen vectors could be applied to the estimation of variance components using Brent method[26], which has fast convergence determined via the absolute tolerance of heritability rather than all variance components; and the results of eigen decomposition could be also used for solving the mixed model equation when MLM is selected for the association tests. The globally efficient calculation design of GWAS process makes rMVP only need to do the eigen decomposition once instead of doing it multiple times, its results could be directly used in calculations of PC derivation, variance components estimation, and association tests, and the computing time is greatly decreased.
6. **Visualization of results.** High-quality figures are generated to display data information, population structure and GWAS results, including marker density plot, phenotype distribution plot, PCA plot, Manhattan plot, and Q-Q (Quantile-Quantile) plot.

## Results

### Memory-efficient: Efficient memory usage in data loading and parallel computation

Genotype matrices are the biggest datasets for GWAS. In rMVP, genotype data in multiple formats are converted to ‘big.matrix’, which can minimize RAM usage through generating a bridge that facilitates RAM accessing the data on the hard disk instead of loading it to RAM directly as the most software tools do. rMVP achieves this goal by using the ‘bigmemory’ package to build data mirrors that are accessible to RAM, while the actual data remain on the hard drive. In this way, very little RAM capacity is needed for the temporary storage of the data. Once the data mirrors are built, users will never need to re-build them again and the time of loading input data is negligible. When multiple threads are used to accelerate the association tests, no additional data mirrors will be copied for each thread as all threads will share the same data mirrors.

Here, we made a rough illustration of ‘big.matrix’ based memory storage of one and multiple threads for rMVP. The complete GWAS procedure of three methods was recorded for RAM usage test in a Linux server (‘RES’ – ‘SHR’). In this test, the product of genotype data size was measured in standard R matrix format, and ‘theoretical RAM cost’ for multiple threads in ‘fork’ mode is defined as *r* × *c* × *t* × 8 bytes, where *r* and *c* are the number of rows and columns of a matrix respectively, *t* is the number of threads. From the Figure 1, we concluded that, with more threads, rMVP shares variables in RAM among processers and does not require additional memory compared with single thread by the aid of OpenMP (Open Multi-Processing) parallel acceleration. Moreover, by constructing memory-map file for genotype in disk rather than load it all into RAM, rMVP significantly decrease the memory cost, making rMVP pretty promising in process of big data at a PC with limited computing resources.

**Figure 1.**
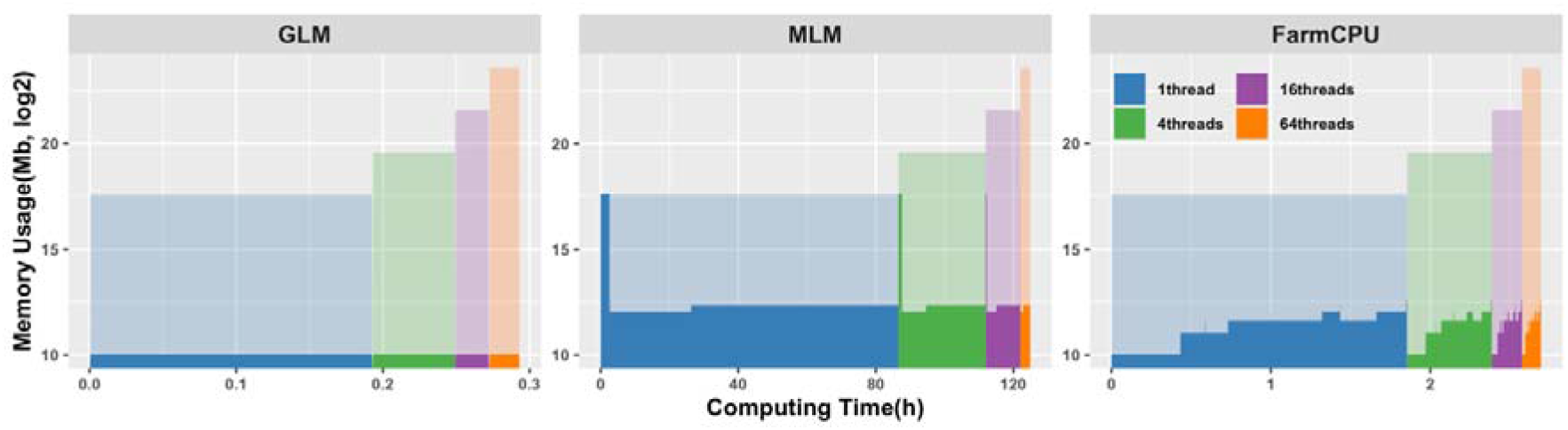
Comparison of memory usage in response to number of threads used for parallel computation under “speed” mode of rMVP. For each block with a specific colour, the y-axis represents memory usage (Mb) in log2 scale; the x-axis represents computing time. Different colour represents different number of threads used for parallel computation, the height of area in dark colour represents real memory costs while the height of shadow in light colour represents theoretical memory costs which is 1, 4, 16, and 64 times of genotype data size in standard R matrix format under ‘fork’ parallel mode, respectively. Data for speed test was generated by PLINK software and each data unit represents 1,000 samples and 100,000 SNPs. The data size for testing memory usage was 16 data units (16,000 samples and 1,600,000 SNPs), 10 PCs are added as covariates in all test methods. All tests were performed on a Red Hat Enterprise Linux sever with 2.60 GHz Intel(R) Xeon(R) 32CPUs E5-4620 v2, and 512 GB memory.

For MLM in Figure 1, a high shoulder peak appears at the beginning of the memory records, which indicating that the most memory cost part of the MLM is the construction of GRM. From the computation details of VanRaden method described above (Equation 4), we can conclude that the calculation of *Z^T^ Z* requires gigantic storage space and the requirement is increasing with both the marker size and the number of individuals. To take care of this problem, we implement two modes (“speed” and “memory”) in rMVP to handle the big data with limited computation resources.

For the “speed” mode, the genotype matrix is stored in R standard matrix format and the transpose of *Z* matrix and the matrix multiplication are carried out by the RcppArmadillo package, which could be automatically speeded up by the Inter MKL math library based on Microsoft R Open platform. However, the big genotype data is loaded into RAM and resulting in a big memory cost as most of the GWAS software tools do, e.g., GEMMA, GCTA, and GAPIT. For the “memory” mode, all the matrices that required for constructing the GRM are stored in the ‘big.matrix’ format and the matrix multiplication of ‘big.matrix’ is implemented by our newly developed C++ function, which could be parallel accelerated by using the OpenMP (Open Multi-Processing) technology. Although it can significantly decrease the cost of memory, a little bit more computing time is required (Table 1). Users can easily adjust the “priority” parameter to get rid of the memory limit or the fastest speed depending on the data size and computing resources.

**Table 1.** Comparison of memory and time cost under modes of “speed” and “memory”.

| Mem (Gb)<br>/Time (min) | Data Units |  |  |  |  |
| --- | --- | --- | --- | --- | --- |
|  | 1 | 2 | 4 | 8 | 16 |
| **Speed mode** | 0.51/0.05 | 3.28/0.15 | 17.80/0.6 | 73.10/3.2 | 285.60/34.70 |
| **Memory mode** | 0.06/0.20 | 0.08/1.61 | 0.17/9 | 0.53/42.12 | 2.06/461.66 |

*Note:* Data for speed test was generated by PLINK software and each data unit represents 1,000 samples and 100,000 SNPs. Parallel computation with 32 CPUs is used to speed up for both two modes. All tests were performed on a Red Hat Enterprise Linux sever with 2.60 GHz Intel(R) Xeon(R) 32CPUs E5-4620 v2, and 512 GB memory.

### Parallel-accelerated: Parallel computation and block matrix multiplication for accelerating association tests

#### Speed Up by Block matrix multiplication

Most GWAS models contain several columns of covariates, such as PCs and Sex, and the linear model function has to be solved for every single tested marker. This process involves the inverse of the design matrix for covariates and the tested markers. Since the covariates were the same for every tested marker, we partitioned the design matrix into sub-matrices according to the covariates and the testing markers. The inverse of the entire design matrix was calculated from the one-time calculation of the inverse of the sub-matrix of covariates. As the number of covariates and markers increased, sub-matrices partitioning significantly saved computing time (Table 2). Block matrix multiplication strategy has been used in all association tests including GLM, MLM, and FarmCPU.

**Table 2.** Speed performance of general linear model with and without using block matrix multiplication strategy.

| Time (Seconds) | Methods |  |
| --- | --- | --- |
|  | Without<br>block matrix multiplication strategy | With<br>block matrix multiplication strategy |
| **0 covariates** | 1,012 | 597 |
| **3 covariates** | 2,853 | 614 |
| **5 covariates** | 4,908 | 623 |
| **10 covariates** | 10,837 | 681 |

*Note:* 0, 3, 5, and 10 covariates are added in both Plink v1.9 and rMVP for testing speed of general linear model without and with block matrix multiplication strategy, respectively. The advantage of block matrix multiplication is increasing when more covariates added as fixed effects. A dataset includes 16,000 samples with 1,600,000 SNPs was generated by PLINK software and used for test. All tests are performed using single thread.

#### Speed Up by Parallel computation

There are two levels of parallel computation implemented in rMVP: Data level parallel (DLP) and Thread level parallel (TLP). For DLP, based on (1) Based on the Microsoft R Open platform, multi-threads have been automatically assigned to speed up the mathematical calculation, such as matrix manipulation. For DLP, association tests on millions of markers are allocated to a group of threads and calculated simultaneously. rMVP switches between the two levels of parallel computation to achieve the highest speed based on the biggest computation requirements in different GWAS procedures. Since three association test methods (GLM, MLM, and FarmCPU) nearly generated the consistent results (Figure S1) and same Power/FDR performance (Figure S2) as related methods in PLINK v1.9 (written in C++, https://www.cog-genomics.org/plink/) and multiple threads version PLINK v2.0 (written in C++, https://www.cog-genomics.org/plink/2.0/), GEMMA (written in C++, https://github.com/genetics-statistics/GEMMA/), and FarmCPU_pkg (R package written in pure R, http://zzlab.net/FarmCPU/), respectively. The rMVP (written in R and C++) was compared with these software packages for speed performance and the computing time was recorded for each software from loading data to generating results files (Figure 2, Table S1). Detailed software version and scripts for computing speed test are provided in Supplementary Table S2.

**Figure 2.**
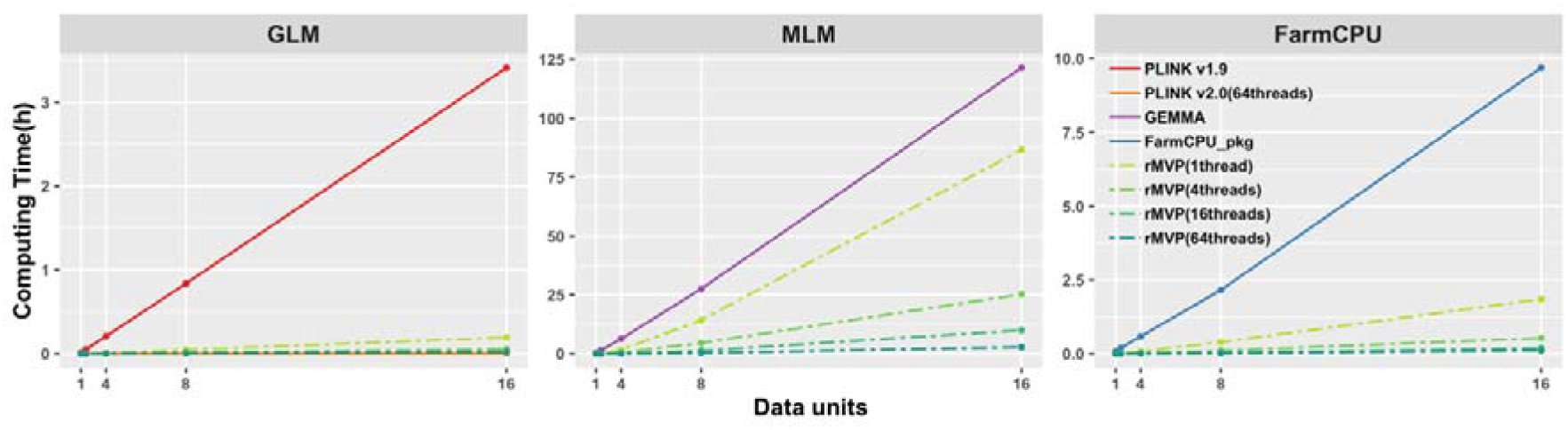
Comparison of computing speed of PLINK, GEMMA, FarmCPU_pkg, and rMVP (“Speed” mode) Computing time (hours) in response to data units are displayed, 5 PCs are added as covariates in all test methods. Speed performances of association test methods in rMVP were performed using 1, 4, 16, 64 threads and compared with PLINK, GEMMA, and FarmCPU_pkg, respectively. Data for speed test was generated by PLINK software and each data unit represents 1,000 samples and 100,000 SNPs. The biggest data for memory test of all models was 16 data units (16,000 samples and 1,600,000 SNPs). All tests were performed on a Red Hat Enterprise Linux sever with 2.60 GHz Intel(R) Xeon(R) 32CPUs E5-4620 v2, and 512 GB memory.

### Visualization enhanced: flexible adjustments for generating high-quality figures

‘MVP.report’ function provides a pack of high-quality figures for visualizing GWAS related information, including data information, population structure, and GWAS results.

Visualization of data information includes phenotype distribution (Figure 3A) and marker density (Figure 3B), which are used to show if the phenotype is normally distributed and the SNPs are evenly covered the entire genome. Skewed phenotype distribution and uneven distributed genotype data would result false positives and biased estimation of population structure and relationship among individuals. Top PCs are visualized in manner of both two and three dimensions to display the population structure (Figure 3G and I).

**Figure 3.**
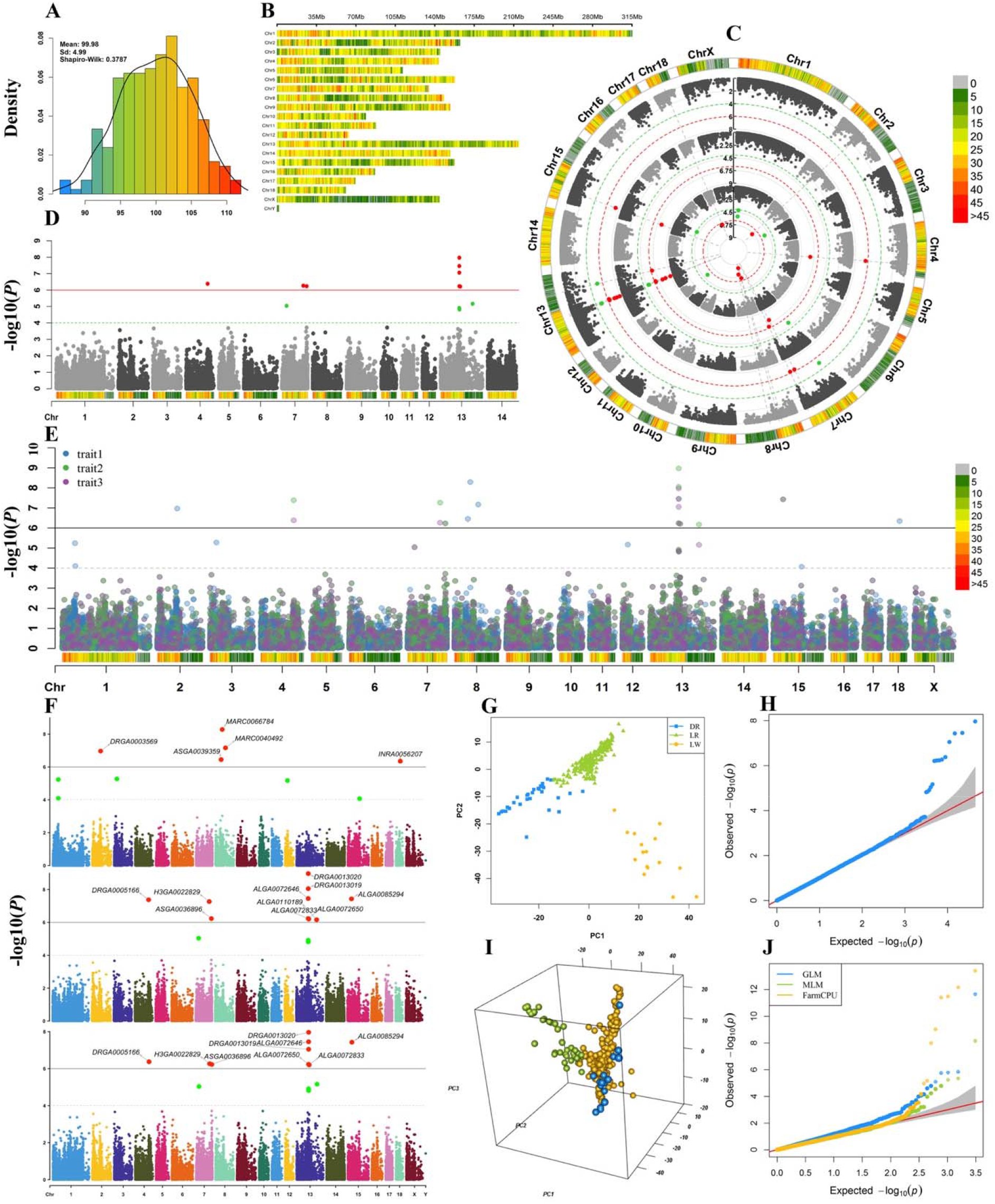
Visualization of GWAS related information. A. Phenotype distribution; B. Marker density, colour lumps with a user-defined window size (e.g. 1 Mb); Manhattan plot for single-group GWAS results with marker density information (D); Manhattan plot for multiple-group GWAS results in both circular manner and rectangular manner (C, E, F); Visualization of population structure in both two dimensions (G) and three dimensions (I); Q-Q plot for single-group GWAS result (H); Q-Q plot for multiple-group GWAS results (J).

Visualization of GWAS results includes Manhattan plot and Q-Q plot. Marker density information is added to Manhattan plot to show the marker coverage of candidate region (Figure 3D). Multiple-group GWAS results can be visualized on a same Manhattan plot and Q-Q plot for easier comparison, common detected signals can be marked with dotted lines, and users could highlight some SNPs or genes of interest on the Manhattan plot without overlap (Figure 3C, E, F, H, and J). Our ‘MVP.report’ can also easily process GWAS results from other software for visualization, such as PLINK, GEMMA, GCTA, and TASSEL. This function can be further extended to visualizing the results from analyses of multi-omics, correlated traits, and eQTL, and to displaying the commonly detected candidate areas. Users can make a desired output figures using more than 40 parameters, detailed description for all parameters are listed in Supplementary Table S3 and Supplementary File S1.

## Discussion

A summary of GWAS related functions of rMVP compared with other software tool is listed in Table 3. At the moment, rMVP does not provide functions of imputation and quality control, which need to be done before association tests. Instead, rMVP provides functions for flexible data conversion that can easily accept the data from other software, such as Beagle[28], which also accepts data in VCF format and provides imputation and quality control functions.

**Table 3.** Summary of GWAS related functions in Plink, GEMMA, FarmCPU_pkg, and rMVP

| Functions | Items | Software |  |  |  |
| --- | --- | --- | --- | --- | --- |
|  |  | Plink | GEMMA | FarmCPU_pkg | rMVP |
| Input | Hapmap | × | × | √ | √ |
|  | VCF | √ | × | × | √ |
|  | Binary | √ | √ | × | √ |
|  | Numeric | × | × | √ | √ |
|  | BIMBAM | × | √ | × | × |
|  | Quality control | √ | × | × | × |
| Model | GLM | √ | √ | √ | √ |
|  | MLM | × | √ | × | √ |
|  | FarmCPU | × | × | √ | √ |
| Population structure | Principal components | √ | × | √ | √ |
|  | Genomic relationship matrix | × | √ | × | √ |
| Variance components estimation | BRENT | × | × | × | √ |
|  | EMMA | × | × | √ | √ |
|  | Fast-LMM | × | × | √ | √ |
|  | HE regression | × | √ | × | √ |
| Output | *p-values*, SE, Effect | √ | √ | √ | √ |
|  | Manhattan plot | × | × | √ | √ |
|  | QQ-plot | × | × | √ | √ |
|  | SNP density plot | × | × | × | √ |
|  | Phenotype distribution | × | × | × | √ |
|  | PCA plot | × | × | √ | √ |

rMVP currently only supports DLP and TLP for parallel computation, lacking the implementation of distributed parallel system (DPS). Compared with TLP that can speed up the computation using 100 threads on a single node, DPS (e.g. MPI, Hadoop, and Spark) can distribute the tasks to 1000 threads on multiple nodes. DPS is also better at dealing with hundreds or thousands of phenotypes and large computing tasks that need to be split, but its performance is limited by the efficiency of data transfer among multi-nodes through the local network. However, association tests in rMVP can be accomplished within 10 hours for a dataset that includes 500,000 samples and five million markers for each sample using FarmCPU model, suggesting that our rMVP can meet most users’ requirements.

Future work includes implementing efficient imputation and quality control functions, and supporting DPS to meet the challenge of big datasets with millions of samples. We also plan to incorporate more association test methods, such as logistic regression and multi-trait model, which fits binary and multi-genetically-correlated traits. With the development of GPU technology, we can get thousands of cores and higher memory bandwidth at a low price. Most of the processes in the GWAS analysis have good independence and can give full play to the advantages of GPU parallel computing. But the bottleneck of limited GPU memory makes it difficult to perform GPU-based GWAS analysis on a large population. In future, we plan to extend rMVP to support parallel computing on multiple machines with multiple GPUs for each machine and explore new memory optimization methods. Incorporating the above methods will greatly improve the versatility of rMVP.

## Code availability

The rMVP package is available on both CRAN (https://cran.r-project.org/web/packages/rMVP) and GitHub (https://github.com/xiaolei-lab/rMVP).

## Authors’ contributions

Xiaolei Liu, Xinyun Li, SZ designed the study, LY, HZ, and Xiaolei Liu wrote the software. ZT, JX, and DY tested the stability of our package using various datasets. ZZ, MZ, XY gave professional suggestions on the experimental design. Xiaolei Liu and LY drafted the manuscript, and ALL authors contributed to finalizing the writing.

## Competing interests

The authors have declared no competing interests.

## Acknowledgements

We thank all rMVP beta version users for giving their valuable feedbacks through GitHub. This work was supported by the National Natural Science Foundation of China [31672391, 31730089, 31702087, 31701144]; the National Key Research and Development Program [2016YFD0101900]; the Fundamental Research Funds for the Central Universities [2662020DKPY007, 2662019PY011]; the National Science Foundation [DBI 1661348]; and the National Swine System Industry Technology System [CARS-35].

## Figure legends

**Table S1.** Computation speed performances of PLINK, GEMMA, and rMVP for five simulated datasets.

**Table S2.** Software versions and codes used in performance tests.

**Table S3.** Parameter details for flexible visualization of GWAS related information.

**Figure S1**

Title: Comparisons of association results between rMVP and related software.

Legend: x-axis is the computed *p-value* in −log10 format for different GWAS models, y-axis is the computed *p-value* in −log10 format of related software for corresponding GWAS model, the experiment was performed on the simulated 16 data units (16,000 samples and 1,600,000 SNPs).

**Figure S2**

Title: Comparisons of power and false positive discovery for different GWAS models between rMVP and related software.

Legend: The experiment was performed using an Arabidopsis dataset, which includes 1178 individuals and 208794 SNPs, the phenotype was simulated by randomly selected 10 QTNs following a normal distribution with mean 0 and variance 0.1, the heritability was 0.5. The final results were the average of 100 replicates.

**Figure S3**

Title: The road mapping of whole GWAS procedures in rMVP.

Legend: K is the Kinship matrix, also known as Genomic relationship matrix (GRM). EigenK represents the eigen decomposition of GRM. PC represents principal components. VC represents variance components.

**File S1.** Demo scripts and figures for visualization in rMVP

## Supplementary material

**Table S1**

**Table S2**

**Table S3**

**Figure S1**

**Figure S2**

**Figure S3**

**File S1**

## References

[1] Visscher PM, Wray NR, Zhang Q, Sklar P, McCarthy MI, Brown MA, et al. 10 Years of GWAS Discovery: Biology, Function, and Translation. Am J Hum Genet 2017;101:5–22.

[2] Yang J, Zaitlen NA, Goddard ME, Visscher PM, Price AL. Advantages and pitfalls in the application of mixed-model association methods. Nat Genet 2014;46:100–6.

[3] Zhang Z, Buckler ES, Casstevens TM, Bradbury PJ. Software engineering the mixed model for genome-wide association studies on large samples. Brief Bioinform 2009;10:664–75.

[4] Pritchard JK, Stephens M, Donnelly P. Inference of population structure using multilocus genotype data. Genetics 2000;155:945–59.

[5] Price AL, Patterson NJ, Plenge RM, Weinblatt ME, Shadick NA, Reich D. Principal components analysis corrects for stratification in genome-wide association studies. Nat Genet 2006;38:904–9.

[6] Purcell S, Neale B, Todd-Brown K, Thomas L, Ferreira MA, Bender D, et al. PLINK: a tool set for whole-genome association and population-based linkage analyses. The American journal of human genetics 2007;81:559–75.

[7] Yu J, Pressoir G, Briggs WH, Vroh Bi I, Yamasaki M, Doebley JF, et al. A unified mixed-model method for association mapping that accounts for multiple levels of relatedness. Nat Genet 2006;38:203–8.

[8] Zhang Z, Ersoz E, Lai C-Q, Todhunter RJ, Tiwari HK, Gore MA, et al. Mixed linear model approach adapted for genome-wide association studies. Nature genetics 2010;42:355.

[9] Lippert C, Listgarten J, Liu Y, Kadie CM, Davidson RI, Heckerman D. FaST linear mixed models for genome-wide association studies. Nature methods 2011;8:833–5.

[10] Segura V, Vilhjálmsson BJ, Platt A, Korte A, Seren Ü, Long Q, et al. An efficient multi-locus mixed-model approach for genome-wide association studies in structured populations. Nature genetics 2012;44:825.

[11] Li M, Liu X, Bradbury P, Yu J, Zhang Y-M, Todhunter RJ, et al. Enrichment of statistical power for genome-wide association studies. BMC biology 2014;12:73.

[12] Bradbury PJ, Zhang Z, Kroon DE, Casstevens TM, Ramdoss Y, Buckler ES. TASSEL: software for association mapping of complex traits in diverse samples. Bioinformatics 2007;23:2633–5.

[13] Lipka AE, Tian F, Wang Q, Peiffer J, Li M, Bradbury PJ, et al. GAPIT: genome association and prediction integrated tool. Bioinformatics 2012;28:2397–9.

[14] Tang Y, Liu X, Wang J, Li M, Wang Q, Tian F, et al. GAPIT version 2: an enhanced integrated tool for genomic association and prediction. The plant genome 2016;9:1–9.

[15] Aulchenko YS, Ripke S, Isaacs A, Van Duijn CM. GenABEL: an R library for genome-wide association analysis. Bioinformatics 2007;23:1294–6.

[16] Kang HM, Sul JH, Service SK, Zaitlen NA, Kong S-y, Freimer NB, et al. Variance component model to account for sample structure in genome-wide association studies. Nature genetics 2010;42:348–54.

[17] Zhou X, Stephens M. Genome-wide efficient mixed-model analysis for association studies. Nat Genet 2012;44:821–4.

[18] Yang J, Lee SH, Goddard ME, Visscher PM. GCTA: a tool for genome-wide complex trait analysis. The American Journal of Human Genetics 2011;88:76–82.

[19] Liu X, Huang M, Fan B, Buckler ES, Zhang Z. Iterative Usage of Fixed and Random Effect Models for Powerful and Efficient Genome-Wide Association Studies. PLoS Genet 2016;12:e1005767.

[20] Yoon S, Nguyen HCT, Yoo YJ, Kim J, Baik B, Kim S, et al. Efficient pathway enrichment and network analysis of GWAS summary data using GSA-SNP2. Nucleic acids research 2018;46:e60–e.

[21] Korte A, Vilhjálmsson BJ, Segura V, Platt A, Long Q, Nordborg M. A mixed-model approach for genome-wide association studies of correlated traits in structured populations. Nature genetics 2012;44:1066–71.

[22] Zhou X, Stephens M. Efficient multivariate linear mixed model algorithms for genome-wide association studies. Nature methods 2014;11:407–9.

[23] Casale FP, Rakitsch B, Lippert C, Stegle O. Efficient set tests for the genetic analysis of correlated traits. Nature methods 2015;12:755–8.

[24] Kane MJ, Emerson J, Weston S. Scalable strategies for computing with massive data. Journal of Statistical Software 2013;55:1–19.

[25] VanRaden PM. Efficient methods to compute genomic predictions. Journal of dairy science 2008;91:4414–23.

[26] Burch BD, Iyer HK. Exact confidence intervals for a variance ratio (or heritability) in a mixed linear model. Biometrics 1997:1318–33.

[27] Zhou X. A unified framework for variance component estimation with summary statistics in genome-wide association studies. The annals of applied statistics 2017;11:2027.

[28] Browning BL, Zhou Y, Browning SR. A one-penny imputed genome from next-generation reference panels. The American Journal of Human Genetics 2018;103:338–48.

